# Several plant self-incompatibility systems may be controlled by atypical receptor-ligand interactions

**DOI:** 10.64898/2026.02.10.705222

**Authors:** Zongcheng Lin, Maurice Bosch, Vernonica E. Franklin-Tong

## Abstract

Self-incompatibility (SI) is the single most important mechanism utilized by flowering plants to avoid self-pollination, thus preventing inbreeding and promoting outcrossing. Many plant SI systems are genetically controlled by a multi-allelic *S*-locus, containing two tightly linked genes that encode the female and male *S*-determinants. When pollen lands on a “self” pistil, interaction between cognate female and male *S*-determinants induces an SI signalling response, resulting in the failure of self-fertilization. Here, we review currently known SI systems that utilize receptor-ligand interactions to control pollen rejection on the stigma surface. Although detailed molecular and cellular information is only known for the SI systems in the Brassicaceae and Papaveraceae, it is apparent that the *S*-determinants of other SI systems (e.g. in the Poaceae and the Convolvulaceae) are likely to also utilize receptor-ligand interactions to prevent self-fertilization. Strikingly, although most of these systems all appear to utilize cysteine rich proteins (CRPs) as ligands to induce an SI response, only one of these receptors is a receptor-like kinase (RLK); the other “receptors” identified to date are proteins of unknown function, which we propose to be atypical receptors (ATRs). Although many of these receptors were identified some time ago, their atypical nature raises many questions, including how they function mechanistically, how they evolved and whether they are found in other plant cell-cell communication systems.

**Significance Statement:** Self-incompatibility involves the precise recognition and rejection of incompatible pollen, often using a receptor-ligand type of interaction between male and female *S*-determinants. In this review we compare several *S*-determinants that appear to function as novel, atypical “receptors” (ATRs), with no kinase- or other distinct domains. We propose that the discovery of these novel “receptors” suggests that further, as yet, unidentified ATRs could be more widely utilized in angiosperms than currently appreciated.

## Introduction

Cell-cell communication is a fundamental biological process for all organisms. This process, vital for almost every aspect of life, operates via direct cellular contact or remote ligand-receptor interactions. Cell-cell communication thus comprises two essential components: the generation of signalling molecules, and their perception as well as transduction by specific receptors. Small protein/peptide ligands are amongst the most important signalling molecules in eukaryotic organisms, with thousands identified (Teyra *et al*., 2020; Xiao *et al*., 2025). Through binding to corresponding receptors on the plasma membrane, they initiate receptor-ligand mediated intercellular communication and signal transduction, thereby participating in various biological processes such as growth, development, and environmental adaptations such as stress and immune responses (Xiao *et al*., 2025). In plants, two major groups of secreted peptide/small protein ligands have been described: post-translationally modified small peptides, and cysteine-rich proteins (CRPs) (Matsubayashi, 2014). CRPs are small secreted proteins, usually <170 amino acids, typically with an even number of at least four conserved cysteines that form pairs of disulfide bonds (Okuda, 2021). The number and arrangement of cysteine residues is used to classify the CRP classes, as this determines their folding pattern. Of the ∼2400 genes encoding secreted peptides/small proteins in Arabidopsis, it has been estimated that at least 825 of them encode CRPs (Hu *et al*., 2021; Ghorbani *et al*., 2015; Silverstein *et al*., 2007). CRPs are expressed in many tissues and have been demonstrated to play vital roles in plant development, reproduction, immunity and abiotic stress responses. It is worth noting that CRPs show highly enriched expression in reproductive tissues (Bircheneder and Dresselhaus, 2016). Recent genetic, biochemical and molecular studies have revealed diverse CRPs that signal through their corresponding receptors. These include defensins (generally eight cysteine residues), pollen coat proteins (PCPs; eight cysteine residues), LUREs (six cysteine residues), Rapid Alkanization Factors (RALFs; four cysteine residues), Epidermal Patterning Factors (EPFs) and EPF-like (EPFL) peptides (six to eight cysteine residues) (Cheung, 2024; Olsson et al., 2019; Wang et al., 2023). However, even in the model plant *Arabidopsis thaliana*, most annotated secreted peptides/small proteins remain orphans whose receptors are unknown, so there remains much to discover in the future.

In animals, it was established by the 1980s that receptor kinases and G protein-coupled receptors (GPCRs) act as receptors for peptide ligands, transmitting intercellular signals (Gschwind et al., 2004; Liu et al., 2024). However, in plants, the presence of rigid cell walls initially led to scepticism regarding the presence of peptide-mediated cell-cell communication. This view changed in the early 1990s when the first receptor-like kinase (RLK), *ZmPK1*, encoding a putative serine/threonine-specific protein kinase was cloned in maize (Walker and Zhang, 1990). Intriguingly, *ZmPK1* had a predicted extracellular domain similar to the *S-*locus glycoprotein (*SLG*) of *Brassica oleracea*, and this led to the identification of the *S-*locus *Receptor Kinase* (*SRK*) whose predicted structure was similar to ZmPK1 (Stein *et al*., 1991). In 1991, an 18-amino-acid peptide named systemin was isolated from tomato leaves and shown to induce plant defence responses (Pearce *et al*., 1991). A few years later, the identification of further plant peptides such as phytosulfokine, which induced cell proliferation (Matsubayashi and Sakagami, 1996), firmly established the critical role of peptides in plant cell-cell communication. In 1996, a Science editorial titled "*Plants, Like Animals, May Make Use of Peptide Signals*" highlighted work from the Nasrallah laboratory at Cornell linking the peptide ligand *S-*locus Cysteine-Rich (SCR), as a good candidate for interacting with the receptor kinase, SRK in the context of their work on self-incompatibility (SI) in the Brassicaceae, though the formal publication of demonstration of this as a receptor-ligand pair came later; see (Marx, 1996). In 2000, CLAVATA1, a RLK, was shown to function as the receptor for the secreted peptide CLAVATA3, jointly regulating the proliferation and differentiation of shoot apical meristem cells (Trotochaud *et al*., 2000). Around the same time, studies formally establishing the SRK-SCR/SP11 as a receptor-ligand pair regulating SI in the Brassicaceae were published (Kachroo *et al*., 2001; Takayama *et al*., 2001). These papers marked the establishment of the research paradigm in which RLKs serve as receptors for secreted peptides to mediate cell-cell communication and signal transduction in plants. Indeed, both the systemin and phytosulfokine receptors were subsequently also determined to be RLKs (Scheer and Ryan, 2002; Matsubayashi *et al*., 2002).

Since then, many RLK-peptide pairs have been discovered (Dievart *et al*., 2020). Receptor-Like Proteins (RLPs), which share a common origin with RLKs but lack a kinase domain, can also function as cell surface receptors through their ectodomains, such as a leucine-rich repeats (LRR), to mediate peptide signalling (Snoeck *et al*., 2023; Ngou *et al*., 2024). RLK/RLP-peptide modules act to regulate growth and development of plant organs (Lalun and Butenko, 2025) and participate in numerous reproductive processes from pollen-stigma recognition, pollen hydration, germination to gamete fusion (Zhu *et al*., 2024; Zhong *et al*., 2025), modulate plant immunity against pathogens (Bender and Zipfel, 2023), and coordinate responses to abiotic stresses such as salinity, drought, and phosphate deficiency (Cheung, 2024; Jose et al., 2020). Most functionally defined RLKs require a co-receptor for high-affinity ligand binding and receptor activation. These co-receptors can be other RLKs, for example the SOMATIC EMBRYOGENESIS RECEPTOR-LIKE KINASE (SERK) and CLV3 INSENSITIVE RECEPTOR KINASE (CIK) subfamily members (Gou and Li, 2020) or glycosylphosphatidylinositol-anchored proteins (GPI-APs), including LORELEI and its homologues LORELEI-like-GPI-anchored protein1 (LLG1), LLG2, and LLG3 (Cheung, 2024). Recent studies have shown that many peptide ligands recruit and induce receptor/co-receptor heterodimerization in the plasma membrane as a common activation mechanism. There is also recent evidence of RLK cleavage, allowing translocation of their intracellular domain to mediate signalling networks (Yu *et al*., 2025). Thus, the identification of RLKs and RLPs as receptors for small protein/peptide ligands in plants in the last few decades has greatly advanced our understanding of receptors, small protein/peptide ligands, and the signalling networks triggered downstream of their interactions; the reader is directed to reviews for further information (Bender & Zipfel, 2023; Cheung et al., 2020). There are also Receptor-like cytoplasmic kinases (RLCKs) which lack extracellular ligand-binding domains and act as integrators of signals downstream of receptor-ligand interactions, but we will not discuss them here; there is a huge amount of information on them, especially in the context of the plant immune response, and the reader is referred to recent reviews (Hailemariam *et al*., 2024; Liang and Zhou, 2018).

SI in plants is a great example of a cell-cell communication system whereby the female pistils recognize and reject “self” (incompatible) pollen to prevent self-fertilisation. This mechanism promotes outcrossing and maintains genetic diversity by preventing inbreeding. It is estimated that SI is present in ∼40% of angiosperm species and has evolved multiple times independently due to fluctuating selective pressures on selfing and outcrossing (Igic *et al*., 2008). This evolutionary history has generated an array of different SI systems (Muñoz-Sanz et al., 2020; Zhang et al., 2024). Here, we focus on homomorphic SI systems, in which flowers are morphologically similar. These SI systems are genetically controlled by one or several multi-allelic *S*-loci (in the grasses, *S*- and *Z*- loci). Each *S*-locus contains tightly linked and co-evolved genes encoding pollen and pistil specific determinants, enabling recognition of “self” or “non-self” pollen. A hallmark feature of these *S*-determinants is that they are extremely polymorphic (often showing only ∼50% amino acid identity between alleles). The large numbers of *S*-haplotypes, which can exceed 40 (Lawrence, 2000), has led to comparisons with the major histocompatibility complex (MHC) in animals because both are highly polymorphic. The polymorphic nature of the *S*-determinants presumably enables different specificities and recognition of different *S*-alleles. To date, there are five distinct homomorphic SI systems which have been genetically and molecularly characterized. The first is the *S*-RNase-based SI system, found in several core eudicot families, including Solanaceae, Plantaginaceae, Rosaceae, Cactaceae, and Rutaceae. For more details of *S*-RNase-based SI, the reader is referred to (Xue, 2025; Zhang et al., 2024). The other four systems: Brassicaceae SI, Papaveraceae SI, Convolvulaceae SI and Poaceae SI, all appear to involve receptor-mediated signalling triggered by secreted peptide ligands (**Table 1**).

**Table 1.**
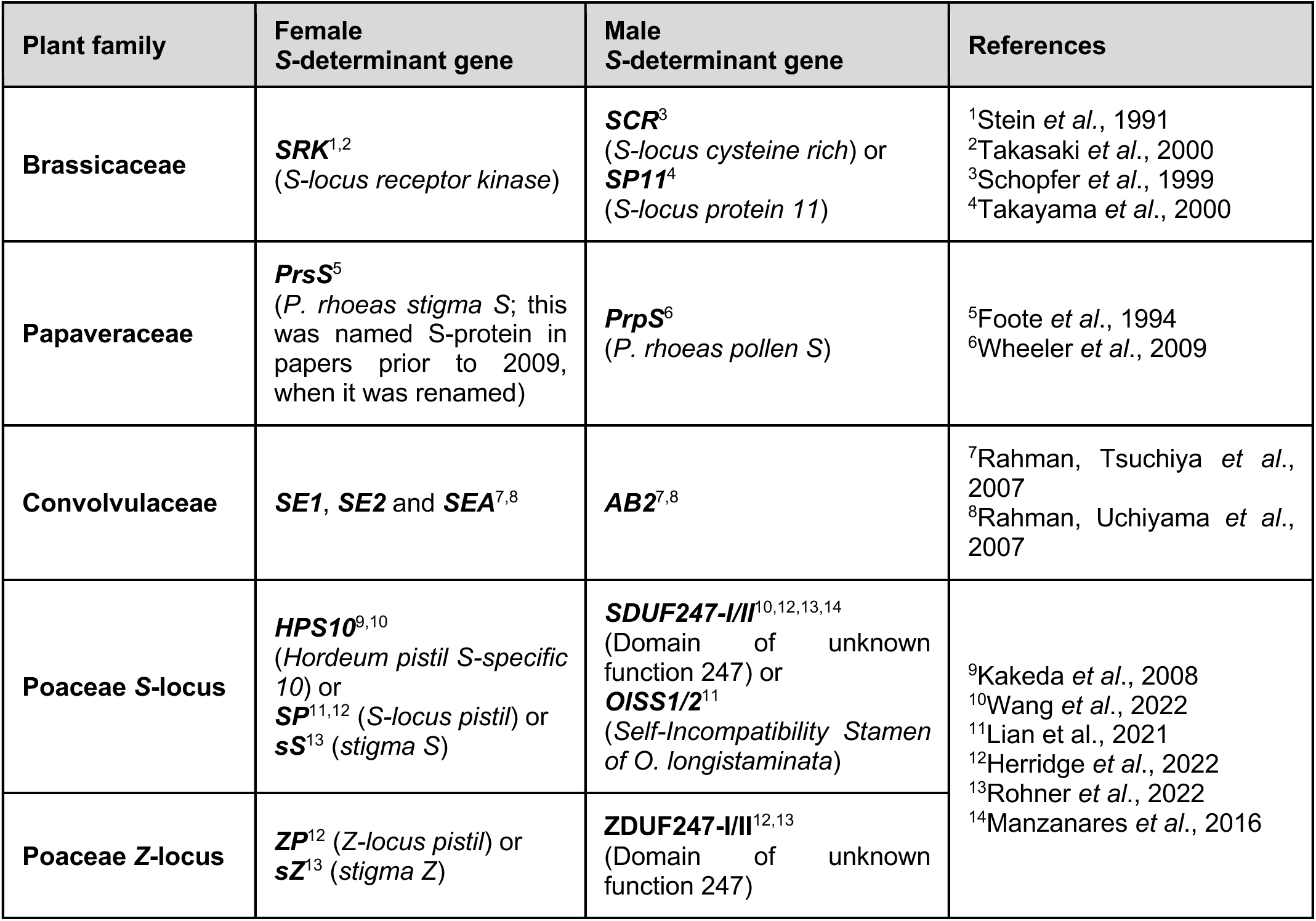
Summary of the *S*-determinants^†^ controlling SI identified to date that are mediated by receptor-ligand signalling. Here we summarize the genes identified at the *S*-/Z-locus as being responsible for controlling SI to date. Because different groups independently identified these genes, in several cases they have been given different names in the literature, which can be confusing to the reader. We have therefore attempted to clarify this in this table, indicating those genes which are homologues in the same box. *^†^For the purposes of clarity, we have called these S-determinants, but as the Poaceae have a 2-locus SI system, they have S- and Z- determinants controlling SI*.

Notably, only the SI system in the Brassicaceae utilizes a classical RLK-peptide ligand pair as *S*-determinants to control SI. In contrast, the *S*-determinants of the other three SI systems (Papaveraceae, Convolvulaceae, and Poaceae) are not RLKs. This suggests the involvement of atypical receptor-ligand interactions to mediate SI in these families. Here we review the *S*-determinants identified in several species that exhibit stigmatic inhibition of incompatible pollen. Although many of these discoveries are not recent, making a comparison here, revisiting them side by side provides a fresh perspective on receptor diversity. We compare the peptide ligands and their (in some cases, putative) receptors identified as *S*-determinants controlling SI and discuss the implications of this in the context of novel receptor systems.

### SI in the Brassicaceae

#### The S-determinants

A sporophytic SI system operates in the Brassicaceae. Incompatible pollen is rapidly inhibited on the stigma surface, usually even before germination (Nasrallah, 2023). The *Brassica* female *S*-determinant, *SRK*, is a member of the large family of plant *RLKs* (Stein *et al*., 1991; Goring and Rothstein, 1992). The cloning of the *SRK* gene at the *S-*locus was inspired by the identification of *ZmPK1*, the first *RLK* found in plants (Stein *et al*., 1991). Subsequently, the *SRK* gene was shown to encode a functional serine/threonine receptor kinase (Goring and Rothstein, 1992) and later, functional transgenic experiments demonstrated that SRK acts as the female *S*-determinant in *Brassica* (Takasaki *et al*., 2000). SRK localizes to the plasma membrane of the stigmatic papillae; it has an extracellular *S*-domain with high homology to a linked *S-Locus Glycoprotein* (*SLG*) gene (which was initially thought to encode the female *S*-determinant (Nasrallah *et al*., 1985), but its function remains equivocal), a single pass transmembrane domain and an intracellular serine/threonine kinase domain (**Figure 1A**). Consistent with a role in SI, the *SRK* alleles from different *S*-haplotypes are highly polymorphic. The soluble extracellular domain of SRK (eSRK) has several conserved domains including twelve conserved cysteine residues and three hypervariable regions (Nasrallah, 2023).

**Figure 1.**
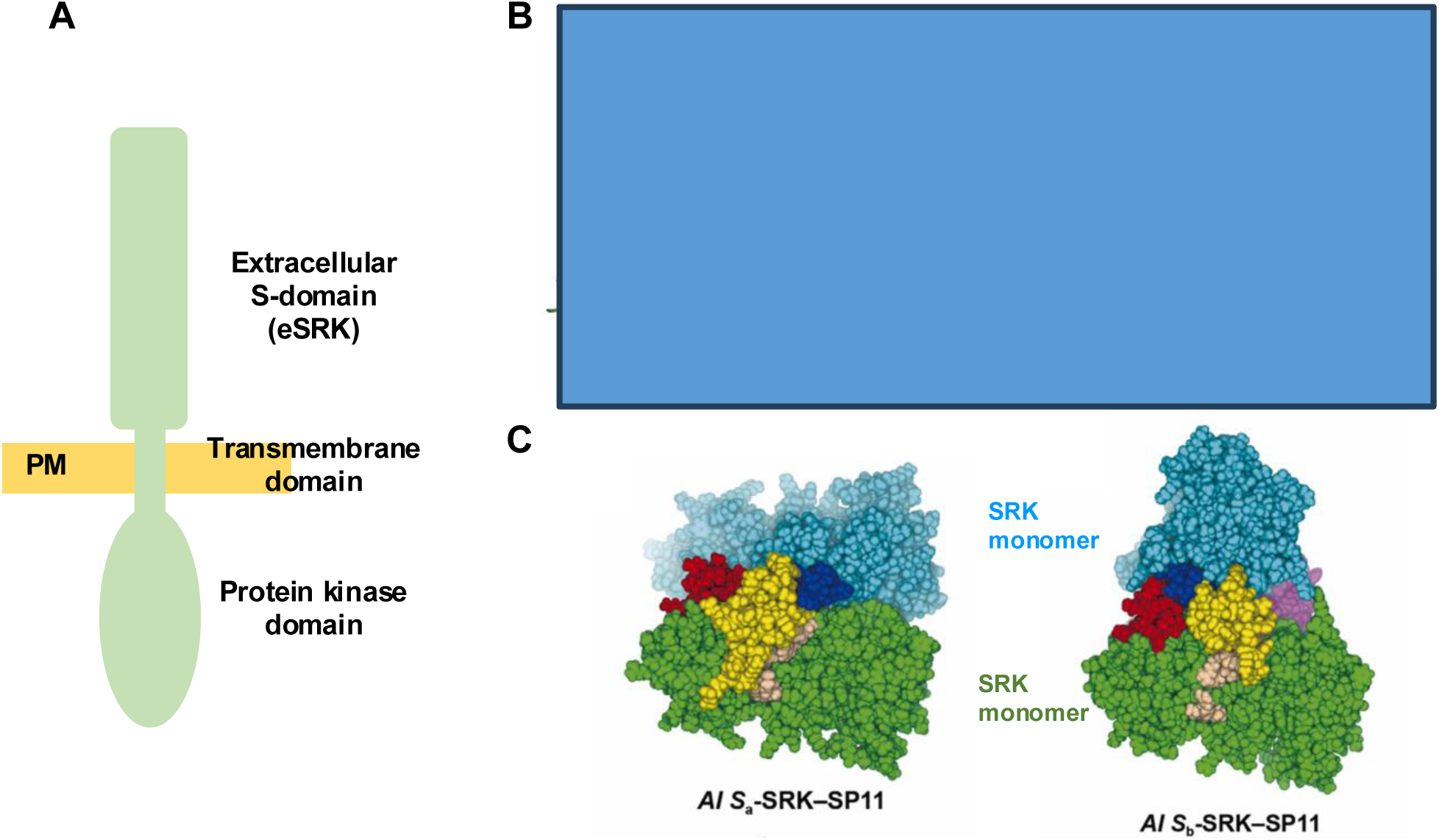
The *S*-locus receptor kinase (SRK) in the Brassicaceae. **(A) Cartoon of the SRK protein.** The *S*-locus receptor kinase (SRK) is a member of the S-domain receptor like kinase (SD-RLK) family. The protein (indicated in green) comprises a highly variable extracellular domain (eSRK), a transmembrane domain and an intracellular serine/threonine protein kinase domain. The eSRK domain has three hypervariable regions that are thought to be involved in generating specificity for the >100 *S*-haplotypes. **(B) Structure of the S_9_-eSRK–SCR/SP11 complex.** Solving the crystal structure for the eSRK has established the interaction of the eSRK domains with cognate SCR/SP11. A cartoon of the secondary structural domains of SRK_9_ is shown above the 3D structures. This indicates the signal peptide (SP), transmembrane domain (TM), two lectin domains (green & cyan), EGF-like domain (yellow), HGF-like domain (purple), twelve conserved cysteine residues and three hypervariable domains (hv1, hv2, hv3). The ribbon structures of the S_9_-eSRK–SCR complex are shown in two orientations. This shows that the S_9_-eSRK dimerizes and binds two S_9_-SCR (purple) to form a heterotetramer (Ma et al 2016). *Copyright permission applied for*. **(C) Predicted binding model for *Arabidopsis lyrata* SRK–SCR/SP11 complexes.** Predicted *A. lyrata S_a_* (left) and *A. lyrata S_b_* (right) complex models using ColabFold. Two SRK monomers are depicted in green and cyan, while two SP11 molecules are shown in yellow and purple. One of the SRK monomers (green) from *S_a_*-SRK and *S_b_*-SRK are aligned in both position and orientation. Three hypervariable (HV) regions of SRK—HV-I, HV-II, and HV-III—are colored pale brown, deep blue, and red, respectively. Among these, HV-I and HV-III belong to the same SRK chain, whereas HV-II belongs to the other chain. Calculations of the binding free energies of the predicted eSRK–SP11 complexes using molecular dynamics simulations showed that some Arabidopsis haplotypes formed a binding mode that was quite different from that of *Brassica rapa S_8_* and *S_9_*. This provided insights into the diversity of haplotype-specific eSRK–SP11 binding modes in Brassicaceae (Sawa et al 2023). *Images taken from Figure 5A-B, Sawa et al (2023) Computational and Structural Biotechnology Journal **21**, 5228-5239. © 2023 The Author(s). Published by Elsevier B.V. on behalf of Research Network of Computational and Structural Biotechnology. (permission not required)*.

The *Brassica* pollen *S*-determinant, *SCR/SP11*, was identified at the *S*-locus almost a decade after the cloning of *SRK* (Schopfer *et al*., 1999; Takayama *et al*., 2000; Suzuki *et al*., 1999) through sequence analysis of the *S*-locus region. The earlier identification of a gene *PCP-A1*, encoding a small highly polymorphic cysteine-rich pollen coat protein (Doughty *et al*., 1998), led to the identification of another protein secreted by the pollen coat, *SP11*, at the *S*-locus and this was proposed to encode the pollen ligand for SRK (Suzuki *et al*., 1999). Independently, the same gene was identified by another group and named *SCR* (Schopfer *et al*., 1999); subsequent transgenic studies showed that SCR/SP11 functioned as the pollen *S*-determinant (Takayama *et al*., 2000; Takayama *et al*., 2001). *SCR/SP11* encodes a small (∼9 kDa), cysteine-rich member of the defensin superfamily, with eight conserved cysteine residues, and polymorphic regions contributing to the *S*-haplotype-specific binding to its cognate SRK (Mishima *et al*., 2003; Murase *et al*., 2020). The first evidence for this came from studies showing allele-specific interaction between SCR/SP11 and SRK in *Brassica* with high affinity, which induced autophosphorylation of SRK kinase domain (Takayama *et al*., 2001; Kachroo *et al*., 2001; Shimosato *et al*., 2007). This provided an unequivocal demonstration that SCR/SP11 acts as a signalling ligand.

The interaction of secreted SCR/SP11 with its cognate SRK at the stigma surface triggers SI in incompatible stigmas. As the eSRK domain is the region critical for interactions with extracellular ligands, structural studies have focused on this domain and have provided insights into the interaction with SCR/SP11 alleles. Studies of the crystal structure of an S_9_-SP11/SCR–S_9_-SRK complex and an engineered S_8_-eSRK in complex with cognate S_8_-SP11 identified critical amino acid residues involved in *S*-haplotype-specific receptor-ligand interactions in *Brassica* (Ma *et al*., 2016; Murase *et al*., 2020). Binding of SCR/SP11 to SRK induces homodimerization of eSRK and the formation of a hetero-tetrameric complex composed of two SRK and two SCR/SP11 molecules, with each SCR/SP11 molecule binding to exposed hypervariable regions of SRK (**Figure 1B**) (Ma *et al*., 2016). These structural studies suggest that ligand recognition between haplotypes differ. Molecular dynamic simulations of self- and nonself-eSRK–SP11 complexes suggest that the *SRK* and *SCR/SP11* genes in *Brassica* have co-evolved to maintain stable interactions between self-combinations and that the *S*-haplotypes can be classified into subgroups with similar recognition modes. The binding free energies are most stable for cognate eSRK-SP11 combinations (Murase et al 2020), and this suggests that this is a feature of this mechanism for self/nonself-discrimination in *Brassica* SI. However, predicting the eSRK–SP11/SCR complex structures for the >100 *S-*haplotypes is challenging due to the high polymorphism of these ligands. More recently, improved structural models for SP11 and the eSRK–SP11 complex have been reported using curated multiple sequence alignments for CRPs to aid modelling of cognate eSRK and SP11 sequence pairs from self-incompatible species of *Arabidopsis* (**Figure 1C**). These models enabled interrogation of the molecular recognition mechanism between cognate pairs at the residue level. Results suggest that further variable regions may contribute to specificity and that the eSRK–SP11 binding mode may vary depending on the *S* haplotypes involved (Sawa *et al*., 2023). No doubt future studies will reveal more about these interactions. Studies also indicated that in *Brassica*, SRK and SLG form a high-affinity receptor complex with SCR/SP11, suggesting some sort of co-receptor function for SLG (Takayama *et al*., 2001). It will be interesting to establish exactly what the role of SLG is and whether it does indeed act as a co-receptor.

### Initiation of SI-triggered interactions at the plasma membrane

Binding of SCR/SP11 to its cognate SRK receptor triggers autophosphorylation of SRK. SRK also interacts with M locus protein kinase (MLPK), which is a receptor-like cytoplasmic kinase (RLCK) localized to the stigma papilla plasma membrane (Kakita *et al*., 2007). Genetic analysis showed that mutation of *MLPK* caused the loss of SI in *Brassica*, demonstrating that MLPK functions as a positive regulator in signalling to mediate SI (Murase *et al*., 2004; Chen *et al*., 2019). MLPK was the first example of a RLCK implicated in mediating RLK signaling; today RLCKs are well established as key players interacting with RLKs to play pivotal roles in mediating signalling networks (Hailemariam *et al*., 2024). More recently, another RLK, FERONIA (FER), has been shown to be implicated in *Brassica rapa* SI by interacting with SRK. During SI pollination, cognate SCR/SP11-SRK interaction enhances the formation of a SRK-FER complex, which triggers increases in reactive oxygen species (ROS) in stigmatic papillae through the activation of FER-regulated NADPH oxidases, causing failure of SI pollen hydration and consequent pollen rejection (Huang et al., 2023; Zhang et al., 2021). However, exactly how SRK-FER-produced ROS triggers inhibition of pollen hydration remains to be established.

### Intracellular events downstream of SRK-SCR/SP11 interaction

One of the earliest detectable physiological outcomes of the SI receptor-ligand interactions initiated at the surface of the stigmatic papilla cell is the inhibition of pollen hydration: activation of the SI pathway blocks water transfer from the stigmatic papilla to the pollen grain within minutes of deposition, so incompatible pollen generally fails to hydrate (Dickinson, 1995; Rozier *et al*., 2020). The signalling network triggered in the stigmatic papilla cells downstream of SRK-SCR/SP11 interaction has been well characterized in *Brassica* and *Arabidopsis*, revealing differences that suggest some redundancy in pollen rejection mechanisms (Goring *et al*., 2023). In transgenic *A. thaliana* expressing *SRK*, self-pollen and chemically synthesized SP11 trigger large increases in cytosolic free Ca²⁺ ([Ca^2+^]_cyt_) in incompatible stigmas; this may be mediated by a glutamate receptor-like (GLR) channel (Iwano *et al*., 2015). Although this was proposed to be a key SI response leading to self-pollen rejection in Arabidopsis, it may not be a universal response; evidence suggests that in *Brassica* this part of the signalling network may have diverged (Goring *et al*., 2023). In *Brassica*, several proteins have been identified as being crucial for SI, interacting with SRK to trigger the signalling network to inhibit incompatible pollen. In addition to MLPK and FER, Thioredoxin h-Like (THL)1/2 also interacts with SRK; this interaction inhibits basal SRK activity (Bower *et al*., 1996; Haffani *et al*., 2004). The U-box E3 ubiquitin ligase ARC1 acts as a key positive regulator of SI downstream of SI signalling initiation, and is phosphorylated by both SRK and MLPK (Gu *et al*., 1998). Activation of ARC1 by phosphorylation results in the ubiquitination and proteolysis of several target proteins, thereby reinforcing SI responses. Three substrates have been identified as targets of ARC1 in *B. napus*: glyoxalase 1 (GLO1) (Sankaranarayanan *et al*., 2015), the EXO70A1 exocyst subunit (Samuel *et al*., 2009), and phospholipase D, PLDα1 (Scandola and Samuel, 2019). These proteins normally promote compatible pollen-stigma interactions. Their ubiquitination and degradation thus disable cellular processes essential for compatible pollen acceptance. For example, the ubiquitination and proteolysis of EXO70A1, triggered by SI in *B. napus*, disrupts exocytosis in the stigmatic papillae, contributing to incompatible pollen rejection. Similarly, degradation of PLDα1, which is also involved in vesicle trafficking and membrane fusion, results in arrest of incompatible pollen. These findings implicate exocytosis and secretion as pivotal control points in both SC and SI responses. Disruption of secretion is thought to underlie the rapid arrest of pollen hydration noted earlier (Zhang et al., 2024), providing a mechanistic link between early physiological observations and the molecular events of the SI pathway. Whether there is a crosstalk between ROS increase and secretion inhibition is an interesting question for future research. The reader is referred to (Goring *et al*., 2023) for more detailed information.

### SI in the Papaveraceae

#### The S-determinants

In contrast to Brassicaceae, in the Papaveraceae the incompatible pollen usually establishes polarity and is either inhibited immediately after this, before the pollen tube emerges or soon after germination (Lawrence, 1975); thus, inhibition is slightly later than observed in Brassicaceae, but it still takes place on the stigma surface. In the Papaveraceae, SI is controlled by a single, multi-allelic *S*-locus, with the pollen phenotype determined gametophytically (Lawrence et al., 1978). Evidence that this SI system involves a receptor-ligand type interaction was established prior to the identification of the *S*-determinants. Addition of stigmatic extracts in an *in vitro* SI bioassay revealed increases in [Ca^2+^]_cyt_ specifically in incompatible pollen tubes (Franklin-Tong et al., 1993, 1997) and was later demonstrated using the recombinant female *S*-determinant (Franklin-Tong et al., 1995). Subsequently, it was shown that *S*-haplotype-specific interaction triggered Ca^2+^ influx involving activation of a non-specific cation channel in incompatible pollen tubes (Wu *et al*., 2011). This triggers a Ca^2+^-dependent signalling network in incompatible pollen, resulting in growth arrest and ultimately programmed cell death (PCD); see later.

The female *S*-determinant was first identified through stigmatic proteins that segregated with the *S_1_* allele, leading to the cloning of the *S_1_* gene, later renamed as *Papaver rhoeas stigma S-determinant* (*PrsS_1_*) (Foote *et al*., 1994). *PrsS* encodes a small (∼15 kDa), secreted, cysteine-rich protein specifically expressed in the stigmatic papilla cells. Sequence information for three *PrsS* alleles in *P. rhoeas* revealed high polymorphism, yet these proteins share a conserved β-strand-rich secondary structure with four conserved cysteine residues that are predicted to form disulfide bonds stabilizing the protein. Despite the high polymorphism, obvious blocks of hypervariable regions are absent; however, site-directed mutagenesis revealed that several amino acids present in three predicted hydrophilic loops (2, 4 and 6) are essential for biological activity (Kakeda *et al*., 1998), so are likely to interact with the male *S*-determinant, *Papaver rhoeas* pollen S (PrpS) (Rajasekar *et al*., 2019). It is thought that loop 6 is most likely to confer specificity, while the exposed part of loop 4 with hydrophobic residues is unusual and is likely to be involved in intermolecular interactions with PrpS (Rajasekar *et al*., 2019). Critically, recombinant PrsS_1_ protein specifically inhibited the growth of pollen carrying the *S_1_* allele in an *in-vitro* bioassay (Foote *et al*., 1994), demonstrating that *PrsS_1_* functions as the female *S*-determinant in *P. rhoeas*. Subsequently, 87 putative PrsS-like sequences were identified from various species within the Papaveraceae; it is likely that some of these are functional *S*-allele sequences, though some may be paralogues (Paape *et al*., 2011). Originally considered an orphan protein, PrsS proteins were later classified within the SPH (S-protein homologue) family (Ride *et al*., 1999), a subset of CRPs. While many CRPs have been shown to be involved in pivotal cell–cell interactions during reproduction (Marshall *et al*., 2011), no function has been identified for any of the SPH proteins to date. The structure of SPH15 was recently solved; it has a β-sandwich structure, comprising eight or nine β-sheets (Rajasekar *et al*., 2019). Intriguingly, this topology is shared with the membrane-binding domain of the toxins pneumolysin and perfringolysin, which form oligomeric rings that form pores in eukaryotic membranes with a large hydrophobic core containing residues from each strand (Rajasekar *et al*., 2019). As PrsS proteins are predicted to share the same topology as SPH15 (Rajasekar *et al*., 2019), they could potentially form pores themselves; if they did so, this might provide a mechanism for the ion influx stimulated by SI.

The male *S*-determinant, *PrpS*, was identified by analysis of a cosmid clone containing *PrsS_1_*, exploiting the tight genetic linkage of the male and female *S*-determinants. *PrpS* is expressed specifically in pollen and encodes a small (∼20 kDa), hydrophobic plasma membrane protein with multiple predicted transmembrane domains (Wheeler *et al*., 2009). Analysis of *PrpS* and *PrsS* alleles indicated they co-evolved; this reinforces the idea that PrpS–PrsS together constitute a tightly coupled receptor–ligand module with mutual dependency for biological function. However, somewhat surprisingly, *PrpS* lacks canonical functional domains such as a kinase motif, leucine-rich repeats, or extracellular binding domains, and shows no sequence similarity to known RLKs or RLPs commonly used by plant signalling systems and searches of the sequence databases reveal that *PrpS* has no homologues outside the Papaveraceae. Thus, PrpS represents a novel class of plant plasma-membrane located receptor, so should be considered an “atypical” receptor (ATR). Compelling evidence supports its function as a “receptor” in the SI response. Ligand-binding epitopes have been previously identified using peptides based on extracellular domains; a 15-mer peptide corresponding to a predicted 35 amino acid extracellular loop region of PrpS_1_ was found to bind cognate recombinant PrsS_1_ in a concentration-dependent manner. On the basis of structural analysis of SPH15, it has been proposed that the central hydrophobic amino acids of PrpS interact with PrsS’s surface residues of loop 4 and the charged hydrophobic residues within PrpS could interact with the charged amino acids on PrsS’s surface loops 2 and 6 (Rajasekar *et al*., 2019). Moreover, antisense oligonucleotides of *PrpS* prevented SI-induced inhibition of pollen in an *S*-specific manner (Wheeler *et al*., 2009). These data demonstrated that PrpS plays a crucial role in SI-induced *S*-haplotype-specific pollen tube inhibition. This, taken together with the evidence that Ca^2+^ signalling is triggered by *S*-specific interactions, strongly suggests that although PrpS is clearly not a “classical” receptor in structural terms, it represents a novel type of plasma-membrane located ATR that operates without canonical signal-transducing domains.

If PrpS has no kinase domain to transduce a signal upon interaction with PrsS, it most likely relies on its own conformational changes and/or associations with other proteins, such as co-receptors upon ligand binding to mediate Ca^2+^ influx that initiates this SI signalling network. Although the structure of PrpS has not yet been elucidated, it was identified as a predicted “topological homologue” of a calcium channel protein from Drosophila, Flower (FWE) (Wheeler *et al*., 2010), which forms homo-multimeric complexes that function as Ca^2+^-permeable channels involved in presynaptic vesicle endocytosis (Yao *et al*., 2009). Like FWE, PrpS contains multiple conserved acidic residues across its known alleles, which may contribute to ion selectivity and hints at a potential channel-forming role. As SI in *Papaver* is initiated by rapid Ca^2+^ influx involving activation of a non-specific cation channel (Wu *et al*., 2011), a possible scenario that SI might trigger multimerization of PrpS to form or regulate a Ca^2+^-permeable channel has been proposed (Wheeler *et al*., 2010). Although this has yet to be formally established, in support of this idea, subsequent Gene Ontology (GO) predictions using FFPred (Cozzetto *et al*., 2016) suggest that PrpS has a transport-related function; “transport” is the top predicted biological process (probability 0.86) and the top predicted molecular function terms include “substrate-specific transmembrane transported activity” and “ion transmembrane transporter activity”. We have used AlphaFold3 (AF3) (Abramson *et al*., 2024) to model possible predicted structures for PrpS_1_. In the presence of Ca^2+^, a trimeric form is predicted with borderline confidence (ipTM = 0.59), with three Ca^2+^ ions predicted at the centre of the trimer (**Figure 2A**). Modelling higher order multimers under the same conditions revealed predictions of an intriguing ring-like structure with Ca^2+^-binding sites lining a pore (**Figure 2B**). These predicted structures provide a framework for considering how PrpS oligomerisation might occur and how this might contribute to SI signalling. While these structural predictions clearly require experimental validation, they are consistent with the possibility that PrpS may be involved in ion transport and are in line with experimental evidence that SI triggers Ca^2+^-influx and increases in [Ca^2+^]_cyt_. Together, the gene ontology and structural modelling predictions support the view that PrpS acts as a novel atypical receptor (ATR); further studies are required to establish its exact nature and function.

**Figure 2.**
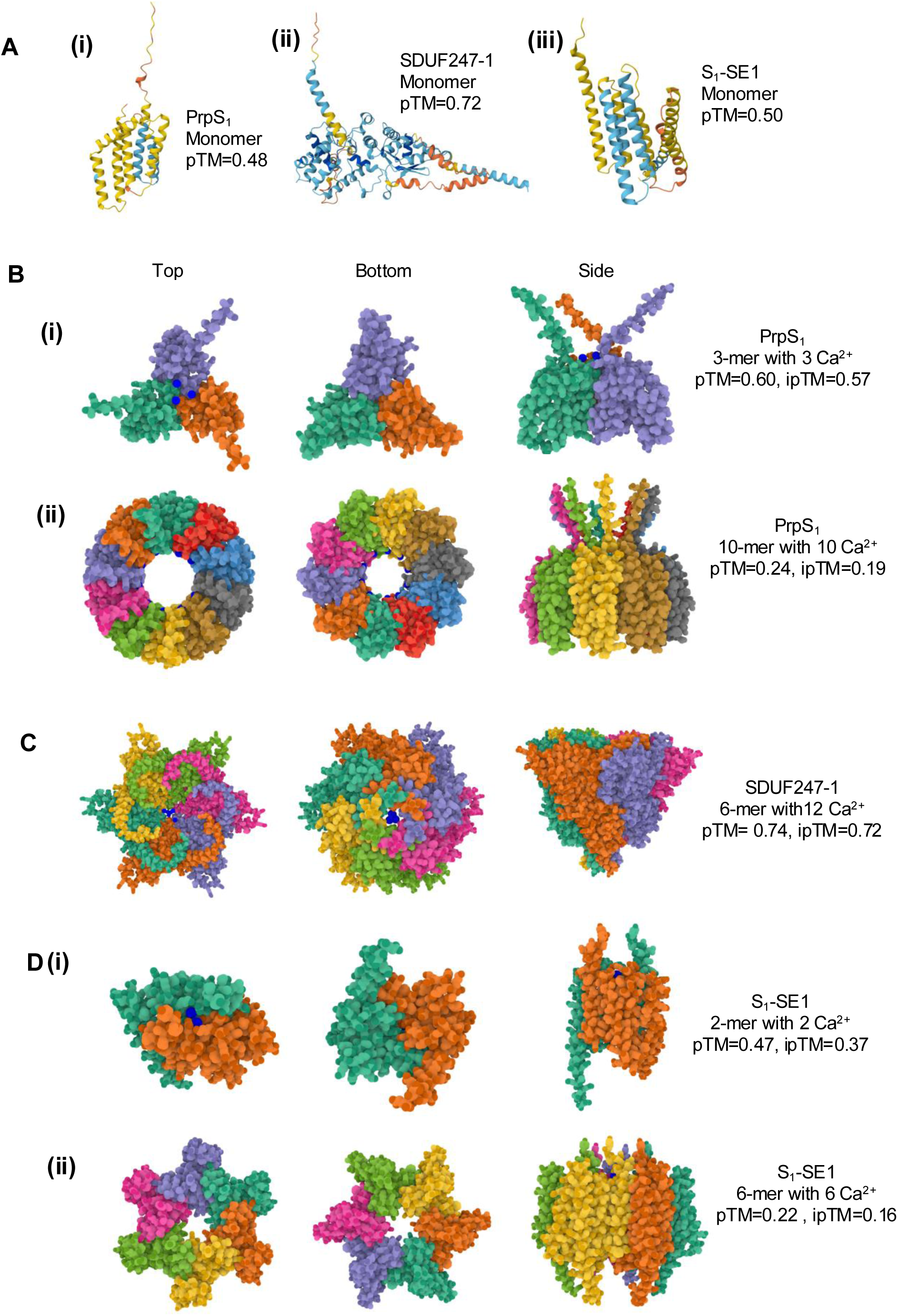
Predictions of monomer (A) and oligomer (B-D) structures for PrpS_1_, SDUF247-1 and S_1_-SE1 proteins using AlphaFold3 (AF3) (*Abramson, J., Adler, J., Dunger, J. et al. (2024) Accurate structure prediction of biomolecular interactions with AlphaFold 3. Nature 630, 493–500*). Oligomeric states were explored systematically by modelling a range of subunit numbers (n = 2 to 10, depending on the protein) in the presence or absence of Ca²⁺. Here we show models corresponding either to the oligomeric state yielding the highest interface predicted template modelling (ipTM) score under the tested conditions, or, in some cases to illustrate possible higher-order architectures despite very low confidence scores. It should be emphasized that these AF3 predictions are presented as conceptual, putative models; no biochemical or biophysical data are currently available to support any of the oligomeric states shown, so these models should be considered highly speculative at this stage. (A) **Predicted monomer structures for (i) PrpS_1_ from *Papaver rhoeas*, (ii) SDUF247-1 from *Lolium perenne*, and (iii) S_1_-SE1 from *Ipomoea trifida* generated by AF3.** The predicted template modelling (pTM) scores for PrpS_1_, SDUF247-1 and S_1_-SE1 monomers are 0.48, 0.72 and 0.50, respectively. The low confidence scores are likely due to the novelty of these proteins, with no homologues identified in available databases. (B) **AF3 predictions for PrpS_1_ oligomer structures.** Oligomers were modelled in the presence of varying Ca^2+^ stoichiometries; predictions were similar in the absence and presence of Ca^2+^ so we only show those in the presence of Ca^2+^. Among the oligomers modelled (n = 2, 3, 4 and 10), a trimer was predicted with the highest confidence: (i) Predicted PrpS_1_ trimer in the presence of Ca^2+^ has an interface predicted template modelling (ipTM) score of 0.57. (ii) Although the pTM (0.19) and ipTM (0.24) scores are extremely low, modelling of higher-order oligomers (n = 10) revealed an intriguing ring-like structure with Ca^2+^-binding sites lining a central cavity. Dark blue dots indicate Ca^2+^ ions. (C) **AF3 prediction for SDUF247-1 oligomer structures.** Oligomers comprising 2-6 subunits were modelled in the presence of varying Ca^2+^ stoichiometries. Here we show a predicted hexamer in the presence of Ca^2+^, which yielded the highest confidence predictions (pTM, 0.74; ipTM 0.72). This displayed a channel-like arrangement with Ca^2+^-binding sites located within a central cavity. Dark blue dots indicate Ca^2+^ ions. (D) **AF3 prediction for S_1_-SE1 oligomer structures.** Oligomers were modelled in the presence of varying Ca^2+^ stoichiometries. Although it should be noted that pTM and ipTM scores for S_1_-SE1 predictions are all very low, a dimer in the presence of Ca^2+^ (i) was predicted with the highest confidence (pTM=0.47, ipTM=0.37). (ii) Intriguingly, a predicted hexamer of S_1_-SE1 proteins, although with extremely low confidence (pTM=0.22, ipTM=0.16), adopts a pore-like structure. Dark blue dots indicate Ca^2+^ ions.

### Evidence for a GPI-AP required for SI suggests involvement of a co-receptor

Further insights into components required for SI came from a forward genetic screen using *A. thaliana* lines expressing the *Papaver* SI system, in which functional *PrpS* and *PrsS* transgenes were introduced to reconstitute the SI response (Lin *et al*., 2015). This screen identified an orthologue of the mammalian GPI-inositol deacylase, PGAP1, named HLD1, as being required for SI in *Papaver* (Lin *et al*., 2022). As PGAP1 is involved in the post-translational remodelling of GPI-APs, this implicates that GPI-APs play a key role in *Papaver* SI. A proposed model suggests that remodelling of as yet unidentified GPI-APs (and also their cleavage/release from the plasma membrane) facilitates their interaction with other SI-related signalling components (Lin *et al*., 2022). As GPI-APs can function as co-receptors, enhancing receptor-ligand interactions by associating with partner RLKs and their CRP ligands (Zhou, 2019; Li *et al*., 2015; Shen *et al*., 2017), this raises the possibility that a GPI-AP(s) co-receptor may be involved in the *Papaver* SI signalling initiation by interacting with PrpS/PrsS and modulate its activity. This intriguing possibility and identification of putative GPI-APs awaits confirmation.

### Intracellular events triggered downstream of PrpS-PrsS interaction

The *Papaver* SI system represents one of the best characterized models for understanding the downstream signalling events that mediate SI. In incompatible pollen, SI triggers rapid arrest of pollen tube tip growth and subsequently a series of events leading to PCD. Here we briefly outline some of the key early signalling events and their targets, but for a detailed understanding of the mechanistic basis underlying SI in this system, the reader is referred to (Goring *et al*., 2023) for recent review. PrsS acts as an extracellular ligand, triggering ∼immediate Ca^2+^ influx in incompatible pollen tubes. This is the earliest known response, activating increases in cytosolic free calcium [Ca^2+^]_cyt_ and a Ca^2+^-dependent signalling network downstream that is responsible for rapidly inhibiting pollen tube tip growth and initiating PCD to ensure permanent rejection of incompatible pollen tubes (Bosch & Franklin-Tong, 2007; Thomas & Franklin-Tong, 2004). An early target of the Ca^2+^ signals is p26, a soluble pyrophosphatase (sPPase) that is phosphorylated in a Ca^2+^/CaM-dependent manner, leading to its inhibition (Rudd *et al*., 1996; Eaves *et al*., 2017; de Graaf *et al*., 2006). The targeting of such a key biosynthetic enzyme through phosphorylation provided the first indication that SI signals may act as a potential master regulatory mechanism to modulate metabolism. More recently, it has been shown that SI induces major changes in cellular energy status involving rapid depletion of ATP levels, implicating energy homeostasis as a pivotal factor in SI-triggered events (Wang et al., 2022). SI also induces phosphorylation of a MAPK (Rudd *et al*., 2003) whose activation contributes to signals that lead to the initiation of PCD (Chai *et al*., 2017). A further early pivotal event triggered by SI in incompatible pollen tubes are increases in ROS (Wilkins et al., 2011) that target numerous pollen proteins with irreversible oxidative modifications (Haque *et al*., 2020). More recently, studies using roGFP2-Orp1, a genetically encoded hydrogen peroxide (H_2_O_2_) sensor, combined with measurements of mitochondrial metabolism have revealed that elevated [Ca^2+^]_cyt_ and cytosolic acidification converge to trigger mitochondrial H_2_O_2_ production. This is accompanied by a loss of mitochondrial membrane potential and impairment of electron transport, glycolysis, respiration, and the TCA (tricarboxylic acid cycle) cycle (Wang et al., 2025). This link between increases in ROS and rapid energetic collapse is likely to be important for committing incompatible pollen tubes to PCD. Extensive remodelling of the actin cytoskeleton is another consequence of SI and not only causes inhibition of pollen tube growth, but is another critical event leading to PCD (Thomas et al., 2006). Together the SI signals integrate to act on multiple intracellular targets in incompatible pollen that converge to mediate PCD; see (Wang et al., 2020; Wilkins et al., 2014). These studies provide a detailed picture of the downstream events triggered by this novel PrpS-PrsS receptor–ligand interaction.

### SI in the grasses

In the Poaceae (grass) family pollen tube growth is arrested at or near the stigma surface within minutes of pollination (Heslop-Harrison, 1982; Shivanna *et al*., 1982). This family possesses a two-locus (*S* and *Z*), gametophytic SI system that was first described in the 1950s through classical crossing experiments in *Secale cereale* (rye) and *Phalaris coerulescens* (Hayman, 1956; Lundqvist, 1954), and later confirmed in *Lolium perenne* (Cornish *et al*., 1979). This system rejects self-pollen only when the allelic determinants at both *S* and *Z* loci match between haploid pollen and diploid pistil. This requirement for dual-locus identity suggests a more complex underlying molecular mechanism. However, the molecular identities of the *S* and *Z* determinants remained unknown until recently. Breakthroughs came from genetic and genomic analyses in *L. perenne*. A pollen-expressed gene encoding a protein with a Domain of Unknown Function (DUF247) was found to co-segregate with the *S*-locus across multiple biparental populations. This gene, named *LpSDUF247*, was highly polymorphic in outcrossing genotypes and disrupted in self-compatible species such as *L. temulentum*, suggesting a functional role in SI (Manzanares *et al*., 2016). Long-read assemblies of the *S*-locus revealed a cluster comprising of two SDUF247 genes adjacent to a small, stigma-expressed peptide gene, named *sS* (stigma-*S*). A comparable organisation was found for the *Z*-locus: two pollen-expressed DUF247 genes (*ZDUF247-I* and *II*), tightly linked to a small, stigma-expressed gene encoding, *sZ* (Rohner *et al*., 2022). These dual-locus clusters are now proposed as the molecular basis of SI in grasses.

*SI-DUF247* (indicating both *SDUF247* and *ZDUF247*) genes have an intronless open reading frame encoding a predicted protein of 508-559 amino acids, with a small cytoplasmic domain at the N-terminus followed by a transmembrane domain and a non-cytoplasmic domain at the C-terminus (Rohner *et al*., 2022; Herridge *et al*., 2022). Alignments of *SI-DUF247* sequences revealed two regions of hypervariability in the 56-62 kDa SI-DUF247 predicted protein and it was proposed that these relatively unstructured loops could be involved in ligand recognition, and/or oligomerization (Herridge *et al*., 2022). Expression of SDUF247-GFP fusion proteins from *Oryza longistaminata* in *Nicotiana benthamiana* confirmed their plasma membrane localization (Wang et al., 2022). Thus, the male determinants encoded at both *S*- and *Z*-loci in the grasses appear to represent atypical “receptors” (ATRs).

The putative stigma-expressed *S*-determinant genes in *L. perenne*, *sS* (stigma *S*) and *sZ* (*stigma Z*), which Herridge et al. (2022) named *SP* (*S-locus Pistil*) and *ZP* (*Z-locus Pistil*), respectively, encode predicted proteins of 82 to 122 amino acids (∼ 8-13 kDa); see **Table 1**, but here we will use *sS* and *sZ* for clarity. The predicted sS/sZ proteins share structural similarities (Herridge *et al*., 2022; Zhang *et al*., 2021), including a predicted signal peptide and two conserved cysteines. Signal peptide prediction indicates N-terminal cleavage, resulting in a predicted extracellular protein of ∼65 amino acids (Herridge *et al*., 2022), consistent with a role as ligands for their cognate DUF247 receptors. These predicted protein sequences display extensive allelic polymorphism, show tight genetic linkage to their *S* and *Z* loci, and show evidence of co-evolution with their respective cognate *DUF247* proteins (Herridge *et al*., 2022), supporting the idea that they have evolved in tandem in order to maintain functional specificity. Taken together, these features strongly support their identity as the female determinants of SI in grasses.

Functional validation of the *DUF247-sS/sZ* model emerged from work in *Oryza longistaminata*, a wild African rice species with a functional SI system (Lian et al., 2021; Wang et al., 2022). Earlier conceptual support arose from studies in *Hordeum bulbosum*, where a stigma-expressed peptide gene, *HPS10* (*Hordeum pistil S-specific 10*), was proposed as the female determinant at the *S*-locus (Kakeda *et al*., 2008; Kakeda, 2009). Notably, *HPS10* is orthologous to both *OlSP* in *O. longistaminata* and *sS/SP* in *L. perenne*, underscoring the evolutionary conservation of the female SI component. The genes *OlSS1* (*Self-Incompatibility Stamen 1 of O. longistaminata*) and *OlSS2* from *O. longistaminata*, encoding pollen-expressed DUF247 proteins, are syntenic and orthologous to their *L. perenne* counterparts (Lian *et al*., 2021). Importantly, CRISPR-Cas9-mediated knockouts in either *OlSS1* or *OlSP* abolished SI, rendering plants fully self-compatible confirming their proposed role in SI (Wang et al., 2022). A direct physical interaction between SDUF247 and sS/SP proteins has been demonstrated using BiFC and SFLC assays, confirming their proposed role in receptor–ligand recognition (Wang et al., 2022). These findings support the role of SDUF247 as a novel, atypical receptor (ATR) and sS/SP as a ligand and indicate conservation of this SI mechanism across divergent grass lineages.

Together, these studies support a working model whereby each locus (*S* and *Z*) encodes a pollen-expressed DUF247 at the plasma membrane and a secreted stigma-expressed sS/SP or sZ/ZP protein. S/Z-DUF247 proteins are proposed to function as atypical receptors (ATRs), and two models have been suggested: (i) separate heterodimers of S- and Z-DUF247s that independently recognise their cognate ligands, with signals from both loci required for rejection (Rohner *et al*., 2022), or (ii) assembly of a S/Z-DUF247 heterotetramer that integrates recognition of both Ss/SP and sZ/ZP to trigger pollen tube growth arrest (Rohner *et al*., 2022; Herridge *et al*., 2022); see **Figure 3**. In both cases, SI is only triggered when both *S* and *Z* determinants match, to be consistent with classical genetic observations (Lundqvist, 1957; Cornish et al., 1979). Structurally, DUF247 proteins lack intracellular kinase domains suggesting that they may require additional partners to transduce signals intracellularly. One possibility is that S/Z-DUF247 forms part of a larger multi-protein complex, potentially interacting with co-receptors that possess kinase activity, analogous to well-established receptor complexes in other plant signalling pathways (DeFalco and Zipfel, 2021). Alternatively, S/Z-DUF247 proteins could potentially oligomerise, like the model proposed for the *Papaver* SI system, where PrpS also lacks established signalling motifs and the mechanism by which the receptor-ligand binding triggers Ca^2+^ influx remains unresolved to date. The FFPred prediction for SDUF247-1 shows transport as the highest probability (0.847 score) for biological process and ion transmembrane transporter activity is one of the top listed functions. Thus, although this protein is completely different from the *Papaver* PrpS, it appears that it is also likely to function in transport-related processes. We have used AF3 (Abramson *et al*., 2024) to model predicted structures potentially formed by SDUF247-1. Interestingly, hexamers in the presence of six Ca^2+^ have the highest iPTM (0.72, **Figure 2C**). The predicted hexamer, although it requires validation, illustrates how oligomerization of SDUF247 might support a transport-related role. Although speculative, such structural models provide a useful conceptual framework for future studies aimed at uncovering the signal transduction mechanisms operating in grass SI. Unequivocal functional evidence confirming that these components represent the complete S- and Z- determinants, and the mechanisms by which the assumed receptor-ligand binding triggers downstream signalling, remain to be resolved.

**Figure 3.**
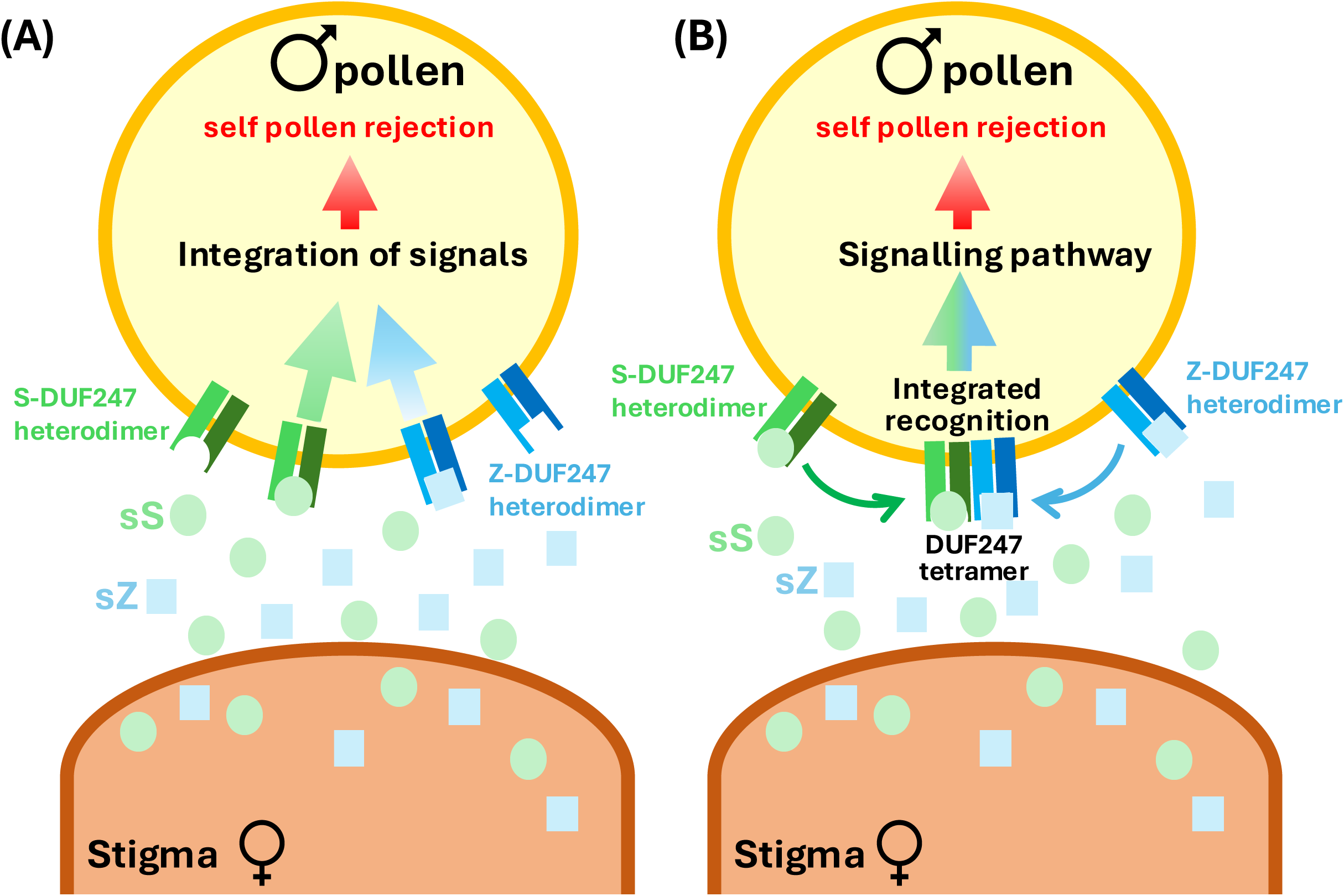
Models for dual-locus recognition and signal integration in grass self-incompatibility (SI) The grass SI system is controlled by two loci: *S* and *Z*. The *sS* and *sZ* genes are expressed in the stigma and encode small proteins predicted to be secreted and proposed to act as signalling ligands. The *SDUF247* and *ZDUF247* genes are expressed in the pollen and predicted to encode transmembrane proteins and are therefore proposed to function as atypical receptors for sS and sZ. To be consistent with classical genetic observations, SI is only triggered when both *S* and *Z* determinants match between pollen and stigma. Two conceptual models have been proposed to explain how S- and Z-specificity might be achieved in the grass SI system (Herridge et al., 2022; Rohner et al., 2023). These two models are illustrated here for an incompatible interaction in which matching S and Z specificities between pollen and stigma result in pollen rejection. **(A) Independent recognition with downstream signal integration.** Pairs of pollen-expressed DUF247 proteins (S-DUF247-I/II and Z-DUF247-I/II, shown in shades of green and blue, respectively) form heterodimers at the pollen plasma membrane. The stigma-expressed ligands sS and sZ (green and blue, respectively) are independently recognised by their cognate DUF247 heterodimers. Ligand binding triggers signalling outputs from each DUF247 complex; these signals are proposed to be integrated downstream, with pollen rejection only occurring when both S- and Z-derived signals are triggered together. **(B) An alternative model proposes that there is integrated recognition through higher-order DUF247 assembly.** Ligand binding to S- and Z-DUF247 heterodimers promotes their assembly into a higher-order S/Z-DUF247 heterotetramer at the pollen plasma membrane. This complex integrates recognition of both sS and sZ ligands, triggering downstream signalling that results in pollen rejection. The molecular nature of the downstream signalling events resulting in incompatible pollen rejection is currently unresolved.

The downstream signalling events triggered in incompatible pollen after the proposed *S-Z* receptor-ligand interactions in the grasses remain largely uncharacterised. Pharmacological studies in rye and *L. perenne* suggest possible involvement of calcium and protein phosphorylation (Wehling *et al*., 1994; Klaas *et al*., 2011), but how these are triggered by DUF247-sS/sZ interaction remains to be established. Nevertheless, SI in the grasses is emerging as an example of a combinatorial receptor-ligand system. Together, these recent advances in knowledge and predictions provide a strong foundation for future studies on receptor activation, downstream signalling and the nature of allelic diversity relating to regulation of SI mechanisms in the Poaceae.

### SI in the Convolvulaceae

*Ipomoea trifida* has a sporophytic SI system determined by a single multiallelic *S*-locus (Kowyama *et al*., 2000), with germination of incompatible pollen inhibited rapidly on the surface of stigmatic papilla, like Brassicaceae. However, studies have shown that *Ipomoea trifida*, a close relative of the sweet potato (*I. batatas*), do not share the same *S*-determinants as Brassicaceae. SI in the Convolvulaceae has therefore clearly evolved independently. To identify the Convolvulaceae *S*-determinants, a positional cloning strategy was adopted. Using linkage mapping, the *S*-locus region was delimited to a 212 kb region in the *S_1_* haplotype (Kowyama *et al*., 2008). The *S*-locus region of the *S_10_* haplotype was identified using similar approaches. Comparison of *S_1_* and S*_10_ S*-locus regions revealed highly divergent regions, named *S* haplotype-specific divergent regions (SDRs), flanked by sequences with high similarities. This result further narrowed the *S*-locus down to the SDR. None of the predicted open reading frames (ORFs) in the SDRs encodes homologues to the currently known SI genes. This confirmed that novel *S*-determinants control SI in the Convolvulaceae. Expression analysis of the genes in the SDRs revealed that three genes, *SE1*, *SE2* and *SEA*, exhibited stigma-specific expression; and one gene, *AB2*, had anther-specific expression (Rahman, Tsuchiya, *et al*., 2007; Rahman, Uchiyama, *et al*., 2007). Moreover, all these four genes exhibited high levels of sequence polymorphism in three *S* haplotypes (Tsuchiya, 2014). This evidence suggests that *SEs* (referring to *SE1*, *SE2* and *SEA*) and *AB2* are strong candidates for the female and male *S*-determinants respectively, in *Ipomoea trifida*.

The anther-specific expressed AB2 proteins are predicted to be small, secreted proteins, comprising 70-80 amino acids, with a putative signal peptide at the N-terminus, and eight conserved cysteine residues at the C-terminus, which are hallmark features of members of the CRP family (see earlier). Thus, AB2 proteins appear to have similarity to the *Brassica* male *S*-determinant, SP11/SCR proteins. These data suggest that, like SCR/SP11, AB2 proteins may function as secreted protein ligands. However, data that it induces SI signalling to mediate inhibition of pollen germination is lacking. Nevertheless, if AB2 acts as a ligand, it implies that there is an AB2 receptor that acts as the female *S*-determinant.

The candidates for the female *S*-determinant in *Ipomoea* are the *SE1*, *SE2*, and *SEA* genes that are expressed in the papilla cells of the stigma. The proteins encoded by the SEs, which are all homologues, are predicted to have four transmembrane domains, but no other functional domains could be identified. The *SE* genes are Convolvulaceae specific, and the predicted proteins show no homology to proteins outside this family in the databases. The predicted structural features of the SE proteins resemble those of the PrpS proteins, the male determinant of SI of *Papaver* (Wheeler *et al*., 2009), and the *Drosophila* FWE protein (Yao *et al*., 2009) that functions as a Ca^2+^ channel. Due to these similarities, it has been hypothesized that SEs might function as the receptors of the AB2 peptide ligand to induce ion signalling in the stigmatic papilla to inhibit germination of pollen (Tsuchiya, 2014). Thus, it appears that the female *S*-determinant in *Ipomoea* is also an atypical receptor (ATR). Using AF3 (Abramson *et al*., 2024) to make structural predictions, hexamers of S1-SE1 proteins appear to look like pore proteins too, although it should be noted that the scores are extremely low, with a high likelihood of being incorrect; this is probably because these are novel proteins (**Figure 2C**). Independent of the structural modelling predictions, FFPred analysis of SE1 predicts involvement of these proteins in transport-related biological processes (probability 0.88) and ion transmembrane transporter activity (probability 0.89), suggesting a potential role in ion transport. However, experimental investigations are required to verify this hypothesis. Nevertheless, like poppy and the grasses, an RLK-type receptor is not used in *Ipomoea trifida* and the evidence strongly suggests that this represents yet another SI system that is likely to utilize a novel atypical receptor (ATR) protein.

### Concluding comments

Plant SI involves a tightly regulated pollen-pistil recognition mechanism and has served as a valuable model system for studying cell-cell communication in plants. Studies of Brassicaceae SI identifying SRK established the research paradigm that RLKs can function as receptors for small protein ligands in plants. Although the SI system in the Brassicaceae utilizes a well-established RLK receptor to interact with its CRP-type ligand, SCR/SP11, it appears that several other SI systems, which all utilize small secreted proteins, mostly members of the CRP family, as their ligands (PrsS in *Papaver*, AB2 in *Ipomoea* and sS/sZ in the grasses), employ novel, atypical receptors (ATRs). What is striking is that, although much remains to be learnt about these novel “receptors”, including their crystal structures and ligand-binding mechanisms, current structural predictions for PrpS and DUF247 (Abramson *et al*., 2024), suggest structural similarities, despite these proteins being otherwise quite distinct, with the possibility that both may have transporter-related or ion channel-like functions. Notably, PrpS, DUF247 and the SE proteins are all highly polymorphic and show no homology to each other or any known receptor proteins, underscoring that these ATRs represent fundamentally distinct molecular solutions to SI-recognition. An important question for the future is how did these proteins evolve? Although *PrpS* and *SE*s are lineage specific genes, only found in the Papaveraceae and Convolvulaceae, respectively, homologues of *DUF247* are ubiquitously found across the angiosperms. It would be interesting to establish whether other *DUF247* gene family members (which are numerous) in other plant families also encode proteins that mediate receptor-ligand signalling.

The discovery of these novel ATRs involved in regulating SI also prompts a broader question: do atypical receptors mediate other signalling pathways mediated by small, secreted proteins in plants in contexts beyond reproductive interactions, for example, in vegetative tissues, during development? The CRP family is huge, and the majority of CRPs are still orphan proteins. De-orphanization of CRPs, and other post-translationally modified small protein ligands, represents a major challenge for future research. This is an important direction for the future, as it will greatly enhance our understanding of fundamental aspects of cell-cell communication in plants, and even beyond. However, although this represents an important open field for future studies, it is technically challenging. Historically, the majority of receptors identified to interact with small protein/peptide ligands belong to the RLK or RLP families, largely because they can be identified by sequence homology. Many researchers to date have used reverse genetic and biochemical approaches by focusing on RLK/RLP family proteins. For example, the receptor for AtLURE1 peptides, PRK6, was identified by screening T-DNA mutant lines for pollen-specific RLK genes (Takeuchi and Higashiyama, 2016). Receptors for Root Meristem Growth Factors (RGFs) were found by using an exhaustive binding assay strategy with a RLK expression library (Shinohara *et al*., 2016). These studies demonstrate that targeting receptor families based on a well-established receptor-ligand paradigm is an effective strategy. Therefore, establishing and validating new modes of receptor-ligand interaction, beyond the classical RLK/RLP paradigm, will be essential for accelerating de-orphanization of small protein and peptide ligands in plants, and bringing new insights about receptor-ligand signalling. How could this be undertaken? We propose the following three approaches:

One approach could involve development of high-throughput, unbiased biochemical methods for receptor identification (“receptor fishing”). Classical interaction assays, including yeast-two-hybrid, coimmunoprecipitation, and photoaffinity-labelled peptides, have been instrumental in validating RLK/RLP-ligand pairs and can be adapted to candidate ATRs. Approaches such as chemical cross-linking, proximity labelling, and ligand-induced proteome profiling offer powerful ways to capture transient or weak receptor-ligand interactions in native cellular contexts.

A second, potentially powerful approach that has recently become available, is the possibility to use Artificial Intelligence (AI)-based approaches as a screening method to identify ATR candidates for peptide ligands. Receptor mediates peptide ligand signalling relies on protein-protein interactions (PPIs) and AI-based prediction of PPIs has significantly enhanced the ability to identify putative ligand-receptor pairs rapidly and cost-effectively compared to traditional experimental methods. For example, AlphaFold2-based approaches have successfully enabled de-orphanization of peptide ligands for single-pass receptors in animal systems (Danneskiold-samsøe *et al*., 2024). Although proteome-wide prediction of PPIs is now feasible (Xiong *et al*., 2025; Zhang *et al*., 2025), current de-orphanization methods still suffer from high false-discovery rates and are limited to well-established receptor families and rely heavily on empirical data. Nevertheless, when combined with genetic, biochemical, and physiological validation, AI-based PPI predictions represent a powerful hypothesis-generating framework for discovering novel ATRs, especially given the rapid development/improvement of AI-based structure and interaction prediction methods.

A third approach could involve establishing novel receptor-ligand paradigm through investigating new biological model systems. For example, SI in angiosperms has evolved independently multiple times, generating a variety of SI systems; it has been estimated that there are at least 35 unique SI systems in flowering plants (Igic *et al*., 2008). As plant SI involves cell-cell recognition and communication, investigating further SI systems has a high potential to identify novel receptor-ligand modules. This has been illustrated in this review by the diverse S-determinants identified to date that utilize ATRs. As the pollen and pistil *S*-determinants controlling SI are tightly linked, advances in genomic research, especially the accurate long-read DNA sequencing technology, provide a unique opportunity to identify pairs of interacting signalling ligands and their corresponding receptor. As it is known that there remain many SI systems that do not correspond to those that are well characterized, this could represent a wealth of information to be mined in the future. Thus, SI systems could serve as valuable models for identification of novel receptor-ligand modules, offering new paradigms for studying cell-cell interactions across diverse biological systems.

The next steps following identification of new ATRs and their corresponding ligands is likely to be a bottleneck, as they will require exhaustive validation and testing of the nature of the putative ATRs experimentally. For example, where structural predictions or sequence features hint at ion channel-like properties or transport-associated functions, electrophysiological approaches, such as heterologous expression followed by patch clamp or planar lipid bilayer recordings, could be used to provide functional validation. This validation stage for each ATR will clearly require significant time, effort and technical investment, but will provide a wealth of information in future years that will expand our knowledge of receptor ligand interactions.

## Acknowledgements

We thank Ms. Wenyue Pei (Huazhong Agricultural University) for AF3 modelling of SI proteins presented in the Figures.

## Funding

Current work in the laboratory of ZL is supported by the National Natural Science Foundation of China (NSFC, grant no. 32270357), and the Foundation of Hubei Hongshan Laboratory (grant no. 11020104). Previous research in the laboratories of MB and VEF-T was supported by the Biotechnology and Biological Sciences Research Council (BBSRC).

## Author Contributions

ZL devised the concept of this manuscript and wrote the first draft. VEF-T and MB contributed significantly to writing and developing the content and theme of this manuscript.

## Conflict of Interest Statement

The authors have no conflicts of interest to declare.

### BOX 1: Main points

- Cysteine-rich proteins (CRPs) function as key signalling molecules for specifying self-incompatibility (SI)
- In addition to the well-characterized S-Receptor Kinase (SRK) involved in Brassica SI, several proteins that function, or are proposed to function, as atypical receptors (ATRs) that have no kinase domain or any distinct domains, have been identified as SI determinants in other species
- The ATRs identified to date are all highly polymorphic and diverse; they share no major sequence similarity, though they share some secondary structure similarities
- Current evidence suggests the possibility that some of these ATRs may function in channel/pore-like roles, although this remains to be demonstrated
- The discovery of these diverse SI determinants suggests that additional, as yet unidentified, ATRs may exist across the plant kingdom

### BOX 2. Open questions remaining

- The discovery of these novel receptors that function as SI determinants suggests that further, as yet unidentified ATRs may exist across the plant kingdom; this could be the “tip of the iceberg”; how many more ATRs will be found in plants?
- How structurally and functionally diverse are these atypical receptors (ATRs) and do they operate through common mechanistic principles?
- What are the molecular mechanisms by which ATRs function?
- Are all ATRs activated by CRPs or are some activated by other classes of ligands?
- How did these ATRs evolve and how have they diversified?

## Notes

### Competing Interest Statement

The authors have declared no competing interest.

